# Exploring Relationships Between *In Vitro* Aqueous Solubility and Permeability and *In Vivo* Fraction Absorbed

**DOI:** 10.1101/2023.11.27.568804

**Authors:** Urban Fagerholm

## Abstract

**Introduction:** Solubility/dissolution and permeability are essential determinants of gastrointestinal absorption of drugs. *In vitro* aqueous solubility (S) and apparent permeability (P_app_) are commonly used as measurements and predictors of *in vivo* fraction absorbed (f_a_) and BCS-classing in humans. The objective of this study was to explore the relationships between *in vitro* aqueous S and Dose number (D_o_) and *in vivo* f_a_ and *in vitro* P_app_ and *in vivo* f_a_ and the predictive power of *in vitro* aqueous S, D_o_ and P_app_.

**Methods:** *In vitro* and *in vivo* data were taken from studies in the literature and correlated. *In vitro* S data were produced in various laboratories and with different methodologies. *In vitro* P_app_ data were produced using Caco-2 and MDCK cells in various laboratories and Caco-2 and RRCK cells in one laboratory each. D_o_ was estimated as oral dose / (S • 250 mL).

**Results:** 452 S data and 1480 P_app_ data were found and used. There was no correlation (R^2^=0.0) between *in vitro* log S and D_o_ *vs in vivo* f_a_, not even at S<1 mg/L or not for compounds with <90 % and <30 % *in vivo* f_a_. A R^2^ of 0.43 was found between log Caco-2 P_app_ and *in vivo* f_a_. The corresponding R^2^ for Caco-2 from one laboratory was 0.65. The interlaboratory R^2^ for the Caco-2 model was 0.48. R^2^-estimates for Caco-2 *vs* MDCK and Caco-2 *vs* RRCK P_app_ were 0.23 and 0.21, respectively.

**Discussion and Conclusion:** Aqueous S appears to have no predictive value of *in vivo* f_a_ in humans, not even at low S or after correction for dose. The shows that one should not base human biopharmaceutical predictions based on aqueous S. Log Caco-2 P_app_ explains about half of the variance of *in vivo* f_a_ in humans. The poor correlations found between Caco-2 and the two other P_app_-models (MDCK and RRCK) demonstrate considerable methodological differences. The unexplained variance does not appear to be explained by S and dose, but rather by *in vitro-in vivo* difference in permeability and poor/absent relationship between *in vitro* S and *in vivo* dissolution potential.

## Introduction

Solubility/dissolution and permeability are essential determinants of gastrointestinal absorption of drugs and the two domains of the Biopharmaceutics Classification System (BCS). *In vitro* solubility (S) and apparent permeability (P_app_), and corresponding *in silico* methods, are commonly used as measurements and predictors of *in vivo* fraction absorbed (f_a_) and BCS-classing in humans.

S is commonly measured using water and buffers as media. Limitations with these are that they do not resemble real human gastrointestinal fluids. According to results by Fagerberg et al. (2015), there is limited consistency between S in phosphate buffer and human intestinal fluid.

Oral dose size can also affect the f_a_. Dose number (D_o_) is a biopharmaceutical parameter that takes both S and dose size into consideration.

Caco-2 is probably the most commonly used cell model for estimation of *in vitro* P_app_ and prediction of *in vivo* f_a_. Other *in vitro* P_app_ cell models include Madin Darby canine kidney (MDCK) and Ralph Russ canine kidney (RRCK) cells.

The objective of this study was to explore the relationships between *in vitro* aqueous S and D_o_ and *in vivo* f_a_ and *in vitro* P_app_ and *in vivo* f_a_, and predictive power of *in vitro* aqueous, D_o_ and P_app_.

## MaterialS & Methods

The literature was searched for studies with large amount of data for *in vitro* aqueous S and P_app_. Useful data were found in the following references: aqueous S, dose size, Caco-2 and MDCK P_app_ from various sources, and *in vivo* f_a_ (Newby et al. 2023; collection of data proprietary of Prosilico), RRCK P_app_ from one source (Keefer et al. 2023) and Caco-2 P_app_ from one source (Thomas et al. 2005).

Simple linear and polynomic methods were used for estimation of correlation coefficients (R^2^) between *in vitro* parameters and *in vivo* f_a_. The method with highest R^2^ was chosen.

D_o_ was estimated as oral dose size / (S • 250 mL).

## Results, DiscussioN & Conclusion

In total, 452 S and 1480 P_app_ (462 Caco-2; 246 MDCK; 678 RRCK, out of which 94 were below the limit of quantification) data were found and used. S-estimates ranged from 0.000097 to 1087000 mg/L and Caco-2 P_app_ ranged from 0.011 to 285 • 10^-6^ cm/s.

There was no correlation (R^2^=0.0) between *in vitro* log aqueous S and *in vivo* f_a_ (n=452) (Figure 1), not even at S<1 mg/L (n=30) or not for compounds with <90 % (n=189) and <30 % (n=30) *in vivo* f_a_, and not between log D_o_ and *in vivo* f_a_.

**Figure 1.**
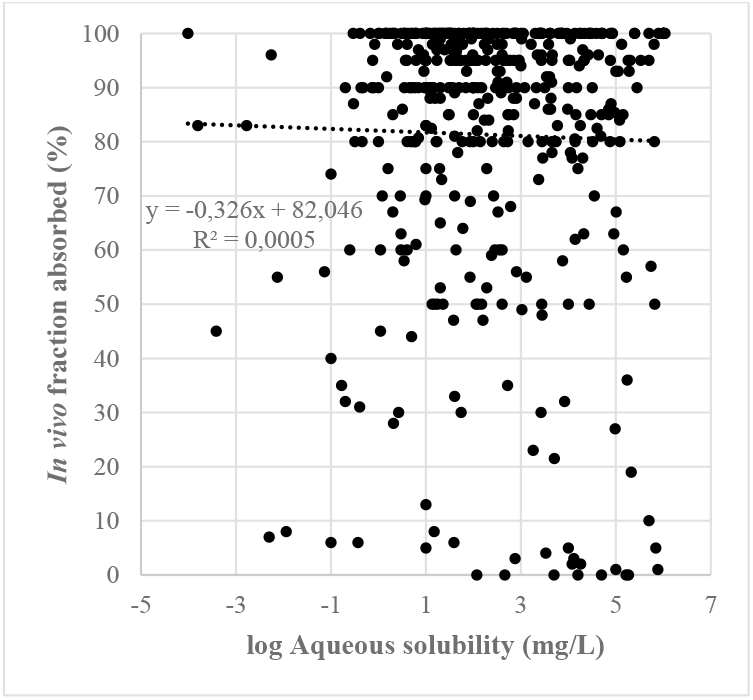
Relationship between *in vitro* log aqueous S and *in vivo* f_a_ (n=452).

A R^2^ of 0.43 was found between log Caco-2 P_app_ and *in vivo* f_a_ (n=380) (Figure 2). For compounds with <90 % the R^2^ was lower, 0.37. The corresponding R^2^ for Caco-2 data from one laboratory was 0.65 (n=79). The interlaboratory R^2^ for Caco-2 was 0.48 (n=64) (Figure 3). At P_app_<5 • 10^-6^ cm/s, however, there was a weak (R^2^=0.13) correlation between laboratories. R^2^-estimates for Caco-2 *vs* MDCK and Caco-2 *vs* RRCK P_app_ were 0.23 (n=185) and 0.21 (n=195), respectively (Figure 4).

**Figure 2.**
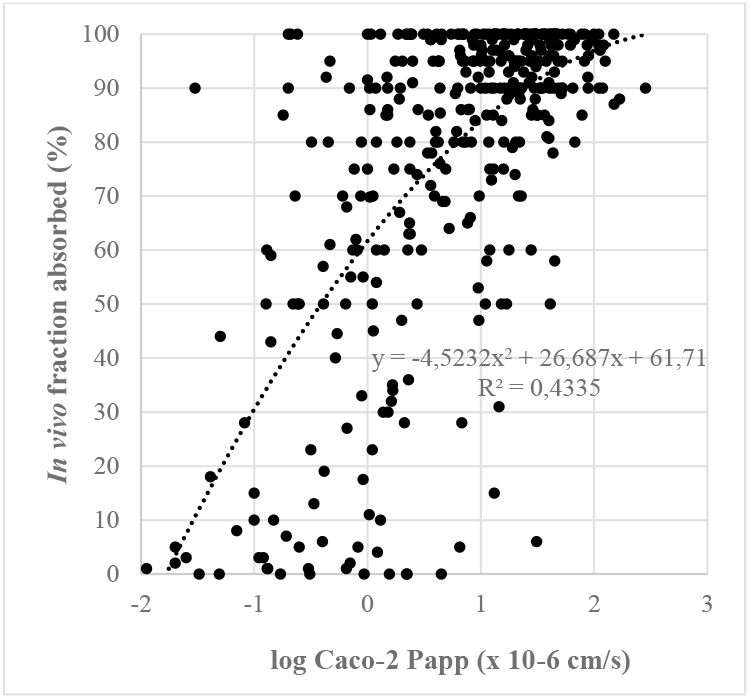
Relationship between *in vitro* log Caco-2 P_app_ (data from various sources) and *in vivo* f_a_ (n=380).

**Figure 3.**
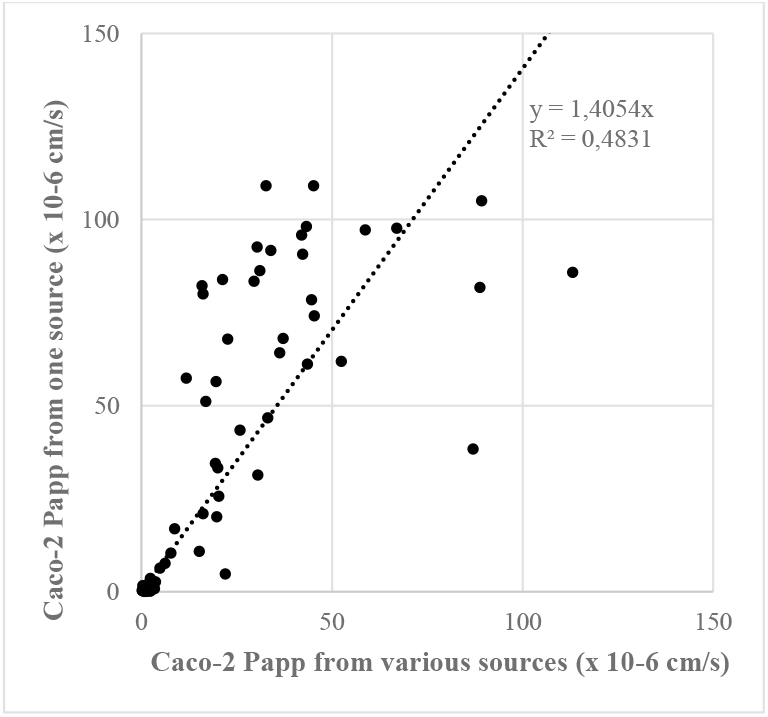
Relationship between *in vitro* log Caco-2 P_app_ from a set with data taken from various sources and from one specific laboratory (n=79).

**Figure 4.**
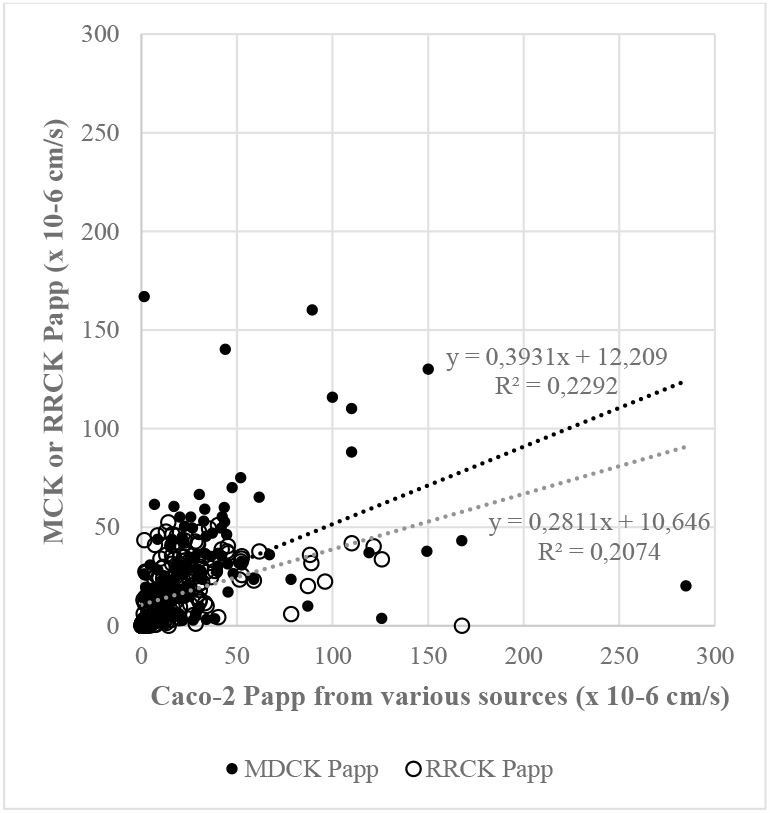
Relationships between *in vitro* log Caco-2 vs MDCK Papp (n=185) and Caco-2 and RRCK P_app_ (n=195).

Complete and near complete gastrointestinal uptake was found for compounds with *in vitro* S, D_o_ and Caco-2 P_app_ of ≥0.000097 mg/L, 4948454 and ≥0.03 • 10^-6^ cm/s, respectively. Less than 10 % *in vivo* f_a_ was found for a compound with *in vitro* S and Caco-2 P_app_ of 39 mg/L and 31 • 10^-6^ cm/s, respectively. For the set with data from various sources, *in vivo* f_a_ ranged from 0 to 100 % over a Caco-2 P_app_ range between 0.2 and 5 • 10^-6^ cm/s. 94 compounds had a RRCK P_app_ below the limit of quantification. The *in vivo* f_a_ ranged and averaged 0-100 and 47 % for 35 of these with available f_a_-estimates, respectively.

Aqueous S appears to have no predictive value of *in vivo* f_a_ in humans, not even after correction for dose size. The shows that one should not base human biopharmaceutical predictions based on aqueous S. Interlaboratory variability is also an obstacle for S. For example, there are 8400- and 4500-fold differences for highest and lowest reported aqueous S for dipyramidole and diclofenac, respectively (Pham-The et al. 2013).

Log Caco-2 P_app_ explains about half of the variance of *in vivo* f_a_ in humans. Higher predictive accuracy can be achieved by avoiding mixing P_app_ data from different sources, at least for compounds with moderate to high P_app_. The poor correlations that were found between Caco-2 and the two other P_app_-models (MDCK and RRCK) demonstrate considerable methodological differences. The unexplained variance does not appear to be explained by S and dose, but rather by *in vitro-in vivo* difference in permeability and poor/absent relationship between *in vitro* S and *in vivo* dissolution potential.

*In vivo* dissolution potential can be predicted using *in silico* prediction methodology. The ANDROMEDA by Prosilico software predicts this parameter with a R^2^ (forward-looking prediction) of 0.57 (n>100; Fagerholm et al. 2022a-c, 2023a,b and unpublished data). This can be compared to R^2^s for log S in water, FaSSIF and human intestinal fluid of 0.00, 0.12 and 0.38, respectively. Preferably, S in human intestinal fluid and *in silico* predicted *in vivo* dissolution potential are used for prediction of *in vivo* f_a_ and BCS in humans.

## References

Fagerberg JH, Karlsson E, Ulander J, Hanisch G, Bergström CAS. 2015. Computational prediction of drug solubility in fasted simulated and aspirated human intestinal fluid. Pharm Res. 32:578–589.

Fagerholm U, Hellberg S, Alvarsson J. Spjuth O. 2022a. In silico predictions of the gastrointestinal uptake of macrocycles in man using conformal prediction methodology. J Pharm Sci. 111;2614–2619.

Fagerholm U, Hellberg S, Alvarsson J, Spjuth O. 2022b. Predicting gastrointestinal absorption of prodrugs and their drugs with the ANDROMEDA by Prosilico software. bioRxiv, Nov 2022.

Fagerholm U, Hellberg S, Alvarsson J, Spjuth O. 2022c. Predicting the influence of fat food intake on the absorption and systemic exposure of modern small drugs using ANDROMEDA by Prosilico software. bioRxiv, Dec 2022.

Fagerholm U, Hellberg S, Alvarsson J. Spjuth O. 2023a. In silico prediction of human clinical pharmacokinetics with ANDROMEDA by Prosilico: Predictions for an established benchmarking data set, a modern small drug data set, and a comparison with laboratory methods. Altern Lab Anim. 51;39–54.

Fagerholm U, Hellberg S, Alvarsson J, Spjuth O. 2023b. ANDROMEDA by Prosilico software successfully predicts human clinical pharmacokinetics of 300 drugs out of reach for in vitro methods. bioRxiv, Nov. 2023.

Keefer CE, Chang G, Di L, Tess DA, Osgood SM, Kapinos B, Racich J, Carlo AA, Balesano A. 2023. The comparison of machine learning and mechanistic in vitro–in vivo extrapolation models for the prediction of human intrinsic clearance. Mol Pharmaceut. 20:5616–5630.

Newby D, Freitas AA, Ghafourian T. 2015. Decision trees to characterise the roles of permeability and solubility on the prediction of oral absorption. Eur J Med Chem. 27:751–65.

Pham-The H, Garrigues T, Bermejo M, González-Álvarez I, Cruz Monteagudo M, Ángel Cabrera-Pérez M. 2013. Provisional classification and in silico study of biopharmaceutical system based on Caco-2 cell permeability and dose number. Mol Pharmaceut. 10:2445–2461.

Thomas S, Brightman F, Gill H, Lee S, Pufong B. 2005. Simulation modelling of human intestinal absorption using Caco-2 permeability and kinetic solubility data for early drug discovery. J Pharm Sci. 48:604–613.

